# Optimal implementation of genomic selection in clone breeding programs - exemplified in potato: II. Effect of selection strategy and cross-selection method on long-term genetic gain

**DOI:** 10.1101/2024.06.21.600034

**Authors:** Po-Ya Wu, Benjamin Stich, Stefanie Hartje, Katja Muders, Vanessa Prigge, Delphine Van Inghelandt

**Affiliations:** Institute of Quantitative Genetics and Genomics of Plants, Heinrich Heine University, 40225 Düsseldorf, Germany; Institute for Breeding Research on Agricultural Crops, Federal Research Centre for Cultivated Plants, 18190 Sanitz, Germany; Cluster of Excellence on Plant Sciences (CEPLAS), Heinrich Heine University, 40225 Düsseldorf, Germany; Max Planck Institute for Plant Breeding Research, 50829 Köln, Germany; Böhm-Nordkartoffel Agrarproduktion GmbH & Co. OHG, 21337 Lüneburg, Germany; NORIKA GmbH, 18190 Sanitz, Germany; SaKa Pflanzenzucht GmbH & Co. KG, 24340 Windeby, Germany

**Author notes:** Corresponding author: Delphine Van Inghelandt.

**Keywords:** Clone breeding, potato, cross-selection method, genomic selection, genetic gain, diversity

## Abstract

Different cross-selection (CS) methods incorporating genomic selection (GS) have been used in diploid species to improve long-term genetic gain and preserve diversity. However, their application to heterozygous and autotetraploid crops such as potato is lacking so far. The objectives of our study were to (i) assess how different CS methods incorporating GS with or without consideration of genetic variability affect both short- and long-term genetic gains compared to strategies using phenotypic selection (PS); (ii) evaluate the changes in genetic variability and the efficiency of converting diversity into genetic gain across different CS methods; and (iii) investigate the interaction effects between different genetic architectures and CS methods on long-term genetic gain. In our simulation results, implementing GS with optimal selection intensities had a higher short- and long-term genetic gain compared to any PS strategy. The CS method considering additive and dominance effects to predict progeny mean based on simulated progenies (MEGV-O) reached the highest long-term genetic gain among the assessed mean-based CS methods. Compared to MEGV-O and usefulness criteria (UC), the linear combination of UC and genome-wide diversity (called EUCD) kept the same level of genetic gain but resulted in a higher diversity and a lower number of fixed QTL. Moreover, EUCD had a relatively high efficiency in converting diversity into genetic gain. However, choosing the most appropriate weight to account for diversity in EUCD depends on the genetic architecture of the target trait and the breeder’s objectives. Our results provide breeders with concrete methods to improve their potato breeding programs.

**Key message:** Cross-selection method considering progeny mean performance and genetic variability can improve long-term genetic gain and keep genetic diversity in clone breeding.

## 1 INTRODUCTION

Potato (*Solanum tuberosum* L.) is one of the most important non-cereal crops for human consumption in the world (http://www.fao.org/faostat/en/). In response to a growing global population, producing sufficient food becomes a big challenge for agriculture (Fŕona et al., 2019). In addition, global crop production is expected to be negatively impacted by climate change due to an increase in extreme temperatures and an alternation of rainfall patterns (Abberton et al., 2016). Thus, developing methods and approaches which increase the efficiency and effectiveness of creating improved and adapted potato varieties is one of the important missions of plant geneticists.

A necessary step to develop varieties requires generating new genetic variability. This can be reached via (1) introducing new alleles, for instance using genetic resource collections (Sanchez et al., 2023); and (2) creating new allelic combinations. The latter happens during meiotic recombinations that occurs after crossing parental genotypes to create segregating populations. Subsequently, superior clones are identified in multi-year testing as variety candidates and new cross combinations are determined to start the next breeding cycle. In a typical clonal breeding program, these steps rely until now mostly on phenotypic records, and take several years. This is especially true in potato because most target traits can only be assessed in the later stages due to the crop’s low multiplication coefficient (Grüneberg et al., 2009), which in turn hampers the increase of genetic gain.

With the advent of genomic selection (GS), genetic gain can be enhanced in both livestock and crop breeding (Alemu et al., 2024). In potato, Wu et al. (2023) have shown via computer simulations that implementing GS into one breeding cycle can improve the short-term genetic gain of the target trait compared to using phenotypic selection (PS). While incorporating GS into breeding programs has been shown to increase long-term genetic gain in diploid crops compared to PS (Gaynor et al., 2017; Gorjanc et al., 2018; Muleta et al., 2019; Lubanga et al., 2022; Sanchez et al., 2023; Werner et al., 2023), the effects of implementing GS on the long-term genetic gain in autotetraploid and heterozygous crops are still unclear. Furthermore, due to the complicated quantitative genetics and the importance of dominance effects in the latter, different trends in implementation of GS can be expected compared to diploid crops, which need to be assessed.

The value of new crosses is commonly predicted by the mid-parental performance based on the phenotypic records of candidate parents (Brown & Caligari, 1989). With the advent of GS, the mid-parental performance can be replaced by the estimated genetic values (EGV) from a trained GS model, which has been shown to improve genetic gain in maize compared to the one based on phenotypes (Allier et al., 2019a; Sanchez et al., 2023). However, as GS is also a truncation selection, it is accompanied by an acceleration of the fixation of favorable alleles. This is because the candidate parents that are intermated for creating the next generation have similar genetic backgrounds, which hinders the generation of new allelic recombinations and limits the long-term improvement of genetic gain (c.f Jannink, 2010). Therefore, maintaining diversity in the breeding populations when selecting new crosses is one possibility to improve long-term genetic gain.

Several studies have proposed different approaches to balance genetic gain and diversity while determining desirable new crosses. Daetwyler et al. (2015) proposed an optimal haploid value to predict the best homozygous line that can be generated from a cross. They showed that this approach can improve genetic gain and preserve genetic diversity better than truncation GS. However, the progenies of a cross in potato are highly heterozygous, and thus, the optimal haploid value is not effective in predicting their phenotypes. Schnell & Utz (1975) proposed the usefulness criterion (UC) to predict the performance of a cross. The UC considers the expected progeny mean (*µ*) and the expected response to selection (*iHσ_G_*) in the first generation progenies: UC = *µ* + *iHσ_G_*, where *σ_G_* is the square root of the progeny variance, *i* the selection intensity, and *H* the square root of the heritability. Using UC to select new crosses has been shown to increase genetic gain compared to mid-parental values in simulation studies on maize populations (Lehermeier et al., 2017; Allier et al., 2019a; Sanchez et al., 2023). Furthermore, Zhong & Jannink (2007) made a modification of the UC, called superior progeny value: S = *µ* + *iσ_G_*. This focuses on progeny mean and variance but ignores heritability. However, depending on the traits, both UC and S can be close to the progeny mean as the variation in progeny mean is much higher than the variation in progeny standard deviation (Zhong & Jannink, 2007; Lado et al., 2017). This aspect limits the advantage of cross-selection (CS) methods like UC and S. Therefore, investigating different weights between progeny mean and progeny variance could affect the efficiency of such CS methods on long-term genetic gain. This, however, has not yet been studied before.

The progeny mean of a bi-parental cross can be predicted by mid-parental performance based on either phenotypic records or EGV from a trained GS model. This can be assessed for inbred populations derived from inbred parents or for hybrids and outbreds in the absence of dominance effects. For diploid species, the progeny mean can be also estimated in the presence of dominance effects (Falconer & Mackay, 1996; Wolfe et al., 2021; Werner et al., 2023). However, no formula is available to estimate the progeny mean for autotetraploid species. Furthermore, it is not easy to obtain a reliable prediction of progeny variance (Mohammadi et al., 2015). With the development of dense genome-wide markers and the advent of GS models, the marker effects can be well estimated (Meuwissen et al., 2001). Recently, several formulae considering linkage disequilibrium and recombination rate in parental lines have been derived to predict the progeny variance (Bonk et al., 2016; Lehermeier et al., 2017; Osthushenrich et al., 2017; Allier et al., 2019b; Wolfe et al., 2021). However, these formulae assume a diploid inheritance and, thus cannot be applied to tetraploid potato.

The simulation of virtual progenies of a cross using a genetic map and phased parental haplotype information is an alternative approach to address the lack of a formula considering autotetraploid inheritance (Bernardo, 2014; Mohammadi et al., 2015). Softwares for this purpose are available (e.g. AlphasimR (Gaynor et al., 2021)) and can be used for simulation in autotetraploid species. The use of the average and variance of EGV among in silico progenies to estimate progeny mean and variance could improve the prediction accuracy of progeny mean compared to mid-parental values and provide a possibility to predict progeny variance for autotetraploid species with heterozygous parents. This aspect, however, has not been examined earlier.

An alternative to UC and the derived methods is optimal cross-selection (OCS) (Gorjanc et al., 2018). The basic idea of OCS is to select a group of bi-parental crosses that maximize the expected progeny mean under a certain constraint of genetic diversity or co-ancestry on the selected population of individuals who serve as parents. Through optimization algorithms (e.g. Kinghorn, 2011), this approach has proven to increase long-term genetic gain in a simulated maize breeding program with a minor penalty on the short-term genetic gain compared to using solely UC (Allier et al., 2019a; Sanchez et al., 2023). However, it is extremely more time-consuming to find the optimal parents and crosses compared to the abovementioned CS methods based on ranking the performance among all possible crosses, especially when many markers and candidates are used in autotetraploid breeding program. This limits its utility, especially for potato breeding.

An alternative possibility to OCS to quantify diversity can be based on the genome-wide variation of a cross itself rather than the variation in the whole population of parents for crosses. This could be measured by the expected heterozygosity (He). Accounting for this element during the selection of new crosses may contribute to long-term genetic gain and simultaneously preserve diversity while being computationally easy to realize. However, to the best of our knowledge, no earlier studies have investigated the performance of such a criterion including the genome-wide diversity of a cross to determine new desirable crosses.

The objectives of this study were to (i) assess how different CS methods incorporating GS with or without consideration of genetic variability affect both short- and long-term genetic gains compared to strategies using PS; (ii) evaluate the changes in genetic variability as well as the efficiency of converting diversity into genetic gain across different CS methods; and (iii) investigate the interaction effects between different genetic architectures and CS methods on the long-term genetic gain in polyploid clone breeding programs.

## 2 MATERIALS AND METHODS

### 2.1 Potato empirical genomic dataset

For this simulation study, a set of 80 tetraploid potato clones genotyped for 49,125 phased sequence variants across 12 chromosomes (Baig et al. in preparation) was randomly selected from a diverse panel of 100 clones. The sequence variants, including single nucleotide polymorphism and small insertion/deletion polymorphisms, have been selected from all possible variants (19,649,193) to have a minor allele frequency *>* 0.05, a missing rate *<* 0.1, and are evenly distributed in windows of 15 kilobases. In addition, their corresponding genetic map information was estimated using a Marey map (for details see Wu et al. 2023).

### 2.2 Breeding programs and selection strategies

This simulation study was based on three main selection strategies in a clonal potato breeding program (Wu et al., 2023): (1) Standard-PS: a scheme following a standard potato breeding program relying exclusively on PS, which serves as benchmark; (2) Optimal-PS: a scheme relying on PS but where the optimal selection intensities during the selection process were determined to maximize genetic gain; (3) Optimal-GS: a scheme based on both PS and GS where the optimal selection intensities during the selection process were determined to maximize genetic gain (Figure S1).

To simulate a long-term potato breeding program, 30 sequential breeding cycles were considered. Each breeding cycle of the breeding program comprised seven stages: cross stage (X), seedling stage (SL), single hills stage (SH), A clone stage (A), B clone stage (B), C clone stage (C), and D clone stage (D). During each breeding cycle, the selection was performed following one of the above described three selection strategies. At the end of one breeding cycle, a defined number of D clones were selected as new parents for the next breeding cycle and intercrossed to create new genetic variation. The details of the approaches used to determine new crosses are described in the next section.

In order to allow a fair comparison of performance across different selection strategies and CS methods, a consistent starting point, called burn-in cycle (C_0_), was required. The procedure of the potato breeding program across 30 cycles is shown in Figure 1 and its details are described in the following:

- Burn-in cycle (C**_0_**)

**–** Step 1: 300 crosses were randomly selected from all possible crosses in the half-diallel among the 80 parents (=3,160, called candidate crosses) and served as a crossing plan. From each cross, the same number of progenies, which were in the following designed as SL progenies, were simulated.
**–** Step 2: Selection processes from SL to D clone stage were conducted according to the chosen selection strategy (Figure S1).
**–** Step 3: The top 20 of the 60 D clones were selected based on phenotypes of the target trait and were, together with the 80 parents of C_0_, considered as candidate parents for cycle 1 (C_1_). Therefore, the number of candidate parents in C_1_ became 100 (i.e. 80+20).
- Cycle 1 (C**_1_**)

**–** Step 1: The performance of all possible crosses in the half-diallel among the 100 parents, excluding the 300 random crosses of C_0_ (= 4,650 candidate crosses), was calculated based on the chosen CS method.
**–** Step 2: Based on the calculated performance from Step 1, the top 300 crosses were selected as the crossing plan and, from each cross, the same number of SL progenies, were simulated.
**–** Step 3: Like Step 2 of C_0_.
**–** Step 4: Like Step 3 of C_0_ except that 20 parents were randomly selected from those candidate parents which were not used in the crossing plan of C_1_, and removed from the candidate parents. Therefore, the number of candidate parents in the next cycle (C_2_) remained 100 (i.e. 80+20-20+20).
- Cycle t (C**_t_**), where t ***>*** 1

**–** Step 1: To (i) mimic the breeder’s approach to keep a reasonable size for candidate parents while focusing on new genotypes, and (ii) reduce computational time, only the candidate crosses which were crosses between the 80 old and 20 new ones and all possible crosses in the half-diallel among the 20 new candidate parents were considered for C_t_ and their performances were calculated according to the CS method.
**–** Step 2 to 4: Like Steps 2 to 4 of C_1_.

**Figure 1:**
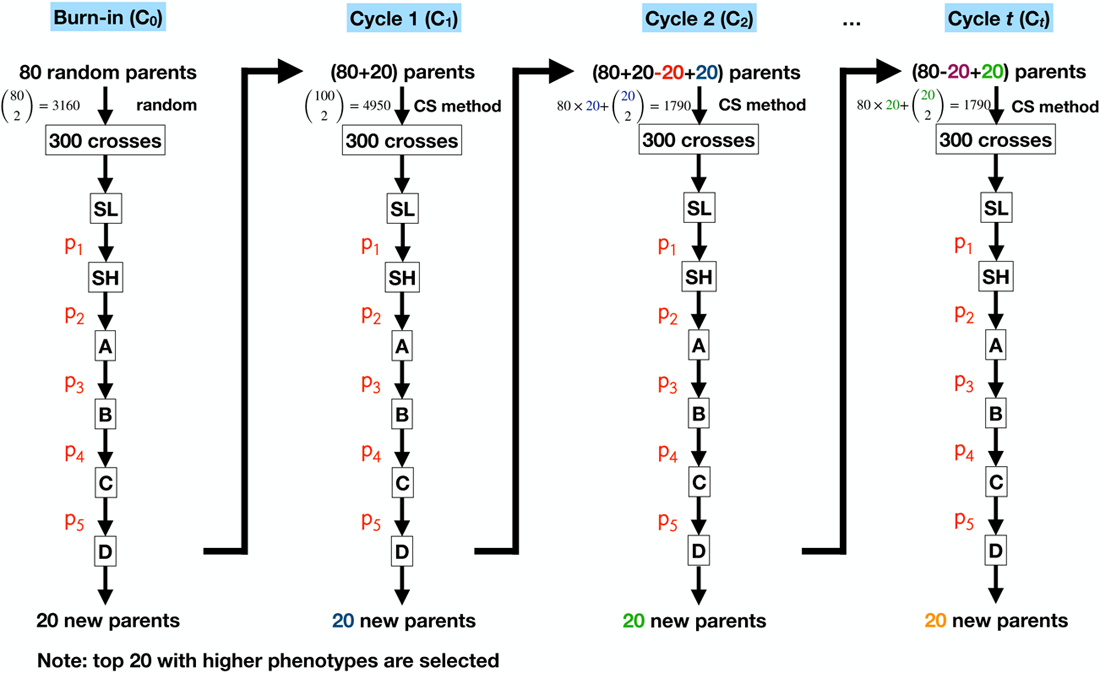
Graphical illustration of recurrent selection in a potato breeding program with the chosen cross-selection (CS) method to determine new crosses. Each breeding cycle of the breeding program comprised seven main stages: cross stage where 300 crosses are selected, seedling stage (SL), single hills stage (SH), A clone stage (A), B clone stage (B), C clone stage (C), and D clone stage (D). p_1_ to p_5_ are selection proportions at each selection stage. Their exact values for each selection strategy and the details about the selection strategies in each breeding cycle are shown in Figure S1 and Wu et al. (2023).

### 2.3 Cross-selection (CS) methods

Different methods were tested to select new crosses for the next cycle. The considered parameters for each cross were (i) the predicted progeny mean, *µ*; (ii) the predicted progeny variance, *σ*^2^_G_; (iii) the predicted progeny diversity; and (iv) the linear combinations of (i), (ii) and (iii).

The predicted progeny mean could be evaluated in five different ways (mean-based CS methods): (i) the mean phenotypic values of the two parents, MPV; (ii) the mean estimated breeding values of the two parents, MEBV-P; (iii) the mean estimated genetic values of the two parents, MEGV-P; (iv) the mean estimated breeding values among simulated offsprings, MEBV-O; and (v) the mean estimated genetic values among simulated offsprings, MEGV-O. The last two, (iv) and (v), were estimated as the mean breeding and genetic values, respectively, among 1,000 simulated progenies of an in silico cross.

To balance the selection of new crosses between improvement of genetic gain and maintenance of variability measured by predicted progeny variance, the concept of UC (Schnell & Utz, 1975) was first extended by:

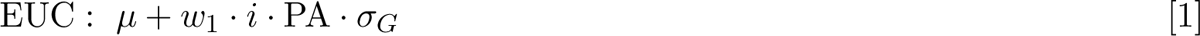

representing an extended usefulness criterion (EUC), in which *µ* was the predicted progeny mean, *w*_1_ a weight on the square root of the progeny variance (*σ_G_*), *i* the selection intensity, and PA the prediction accuracy of the GS model. Here, PA replaced the square root of heritability in the response to selection when GS was implemented (Falconer & Mackay, 1996; Heffner et al., 2010). For EUC, *µ* was based on MEGV-O because this measurement outperformed the other progeny mean estimations in our previous comparison among different mean-based methods (Figure 2a). *σ*^2^_G_ was estimated by the variance of genetic values T_t_ among 1,000 simulated progenies of an in silico cross. *w*_1_ was chosen to be either 1, 10, 50, or 100. If *w*_1_ = 1, the equation [1] is equivalent to UC. Moreover, we assumed the selected proportion per cross as 0.1 so that *i* corresponds to 1.755.

**Figure 2:**
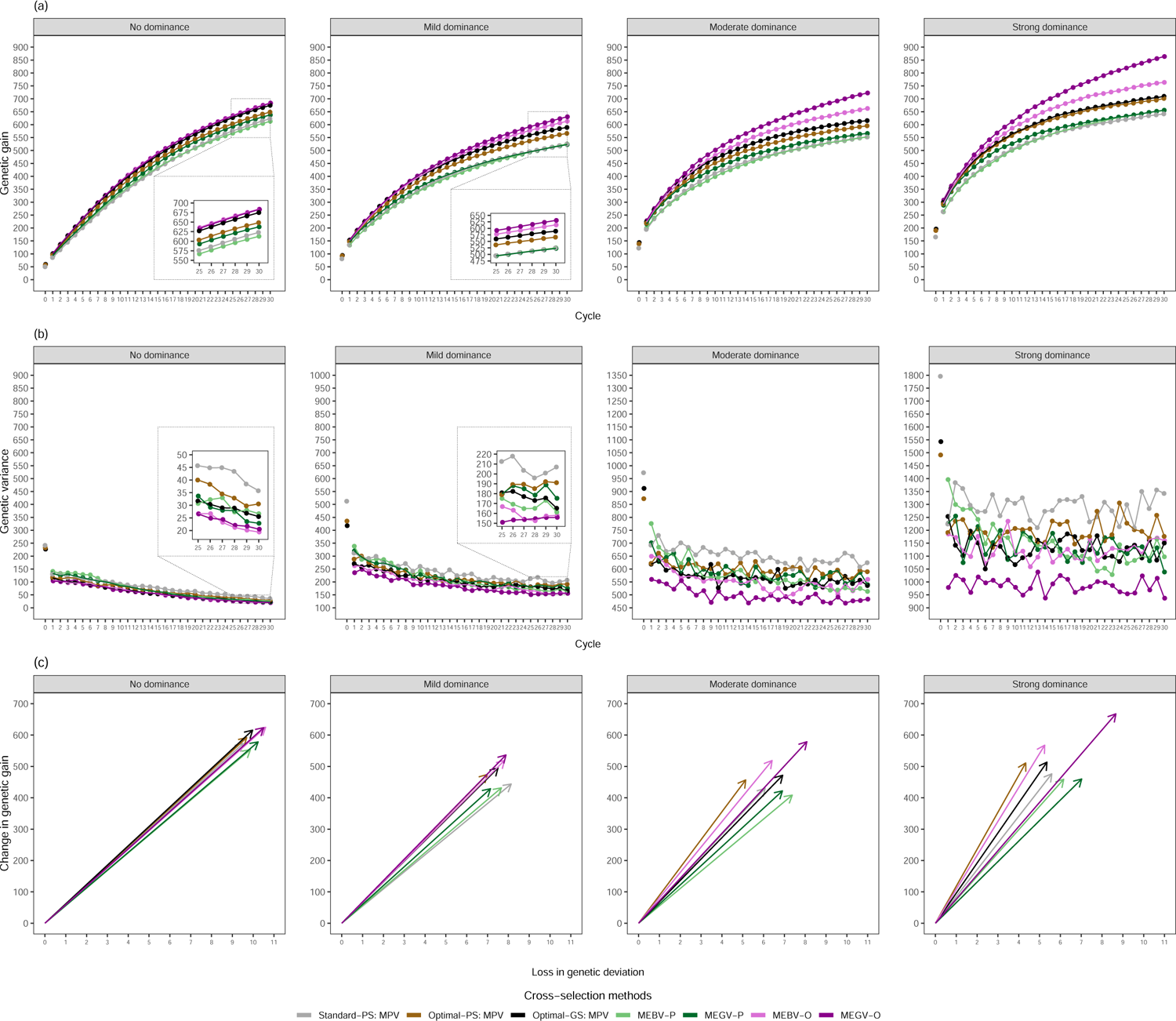
The evolution of genetic gain (a) and genetic variance (b) for the target trait along the 30 breeding cycles on average across 30 simulation runs. The efficiency of converting genetic diversity into genetic gain (c) by regressing the change of genetic gain on the loss of genetic standard deviation between cycle 0 and cycle 30. The three parameters were assessed at D clone stage for different selection strategies (Standard-PS, Optimal-PS, and Optimal-GS), different mean-based cross-selection methods (MPV, MEBV-P, MEGV-P, MEBV-O, and MEGV-O), and different genetic architectures of the target trait (no, mild, moderate, and strong dominance effects).

In addition to EUC and to keep a certain level of genomic diversity in the breeding program, a measure of the gene diversity (as the expected heterozygosity He) was incorporated into the equation [1] to create an extended usefulness criterion incorporating genomic diversity index (EUCD) by:

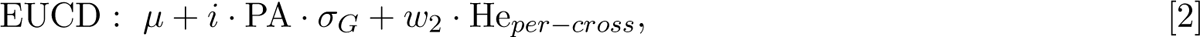

where He*_per−cross_* was used to quantify the genomic diversity of a cross and calculated as the expected heterozygosity He among 1,000 simulated progenies of an in silico cross, and *w*_2_ represented a weight on He*_per−cross_*. Due to the tetraploid nature of potato, He was determined as:

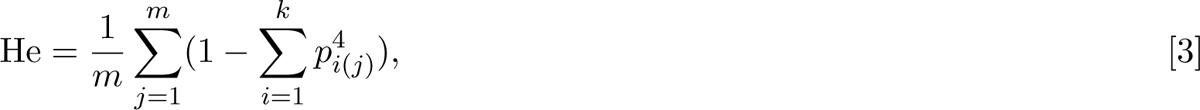

where *m* was the number of sequence variants, *k* the number of alleles in one sequence variant, and *p_i_*_(*j*)_ the allele frequency of the *i^th^* allele at the *j^th^*sequence variant (Gallais, 2003). We only considered bi-allelic sequence variants in this study, and therefore, *k* was equal to 2.

The scale of *σ_G_* and He*_per−cross_* and their variance differed largely. To keep the same level of importance for the two measurements in equations [1] and [2], *w*_2_ was chosen to be 50, 500, 2500, and 5000 (Table 1).

**Table 1:**
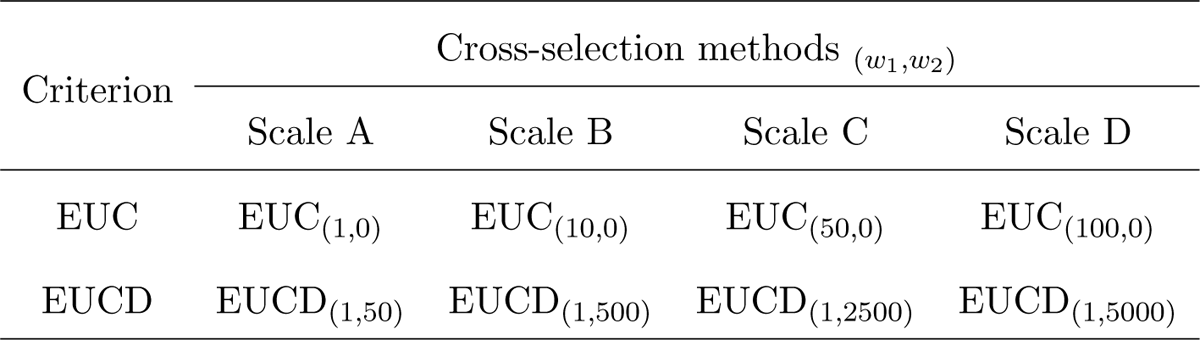
Overview of the different weight (*w*_1_ and *w*_2_) scales for the extended usefulness criterion (EUC) and extended usefulness criterion incorporating genomic diversity index (EUCD), respectively. *w*_1_ is a weight on the square root of the progeny variance, and *w*_2_ a weight on genome-wide diversity quantified by expected heterozygosity (He).

| Criterion | Cross-selection methods $(w_1, w_2)$ | | | |
| --- | --- | --- | --- | --- |
|  | Scale A | Scale B | Scale C | Scale D |
| EUC | $\text{EUC}_{(1,0)}$ | $\text{EUC}_{(10,0)}$ | $\text{EUC}_{(50,0)}$ | $\text{EUC}_{(100,0)}$ |
| EUCD | $\text{EUCD}_{(1,50)}$ | $\text{EUCD}_{(1,500)}$ | $\text{EUCD}_{(1,2500)}$ | $\text{EUCD}_{(1,5000)}$ |

### 2.4 Simulation of genetic architectures of traits

**2.4.1 Simulated true genetic and phenotypic values**

Two traits, auxiliary (T_a_) and target (T_t_) traits, were considered in this study. Here, T_a_ represented the weighted sum of the auxiliary traits measured in the first three stages of the breeding program, and T_t_ the weighted sum of all market-relevant quantitative traits. The latter was controlled by 2,000 quantitative trait loci (QTL). Each QTL included additive and dominance effects, and had five possible genotype classes: aaaa, Aaaa, AAaa, AAAa, and AAAA. The simulation of additive effects across 2,000 QTL followed Wu et al. (2023). The dominance effects, being the deviation of genetic value from the breeding value, were set differently for the three heterozygous genotypes (Aaaa, AAaa, and AAAa) and expressed by d_1_, d_2_, and d_3_, respectively (Gallais, 2003) (Table 2). The simulation of the dominance effects is introduced in the next section. Then, the true genetic value (TGV) for T_t_ was calculated for each clone by summing up the true additive and dominance effects across 2,000 QTL. The TGV for T_a_ was controlled by the genetic correlations between T_a_ and T_t_. The details of the simulated TGV_Ta_ were described in Wu et al. (2023).

**Table 2:** Summary of the five genotype classes, including their coding expression, additive and dominance effects, as well as breeding and genetic values.

| Genotype class | Additive effect (a) | Dominance effect |  |  | Breeding value | Genetic value |
| --- | --- | --- | --- | --- | --- | --- |
|  |  | d<sub>1</sub> | d<sub>2</sub> | d<sub>3</sub> |  |  |
| aaaa | 0 | 0 | 0 | 0 | 0 | 0 |
| Aaaa | 1 | 1 | 0 | 0 | a | a+d<sub>1</sub> |
| AAaa | 2 | 0 | 1 | 0 | 2a | 2a+d<sub>2</sub> |
| AAAa | 3 | 0 | 0 | 1 | 3a | 3a+d<sub>3</sub> |
| AAAA | 4 | 0 | 0 | 0 | 4a | 4a |

The phenotypic values (P) were calculated as P = TGV + *ɛ*, where *ɛ* was a non-genetic value following a normal distribution *N* (0, σ^2^_ɛ_), in which *σ*^2^_ɛ_ was the non-genetic variance. Non-genetic variance for T_t_ (*σ_ɛTt_*^2^) was determined by

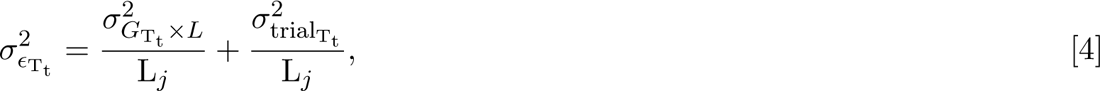

where *σ*^2^_GTt × L_ was the variance of interaction between genotype and location, *σ*^2^_trialTt_ the trial error variance, and L*_j_* the number of location at stage *j*, where *j ∈ {*B, C, D*}* (see Table 1 in Wu et al. 2023). Non-genetic variance for T_a_ (*σ*^2^_Ta_) (= trial error variance, *σ*^2^_trialTa_) was determined by

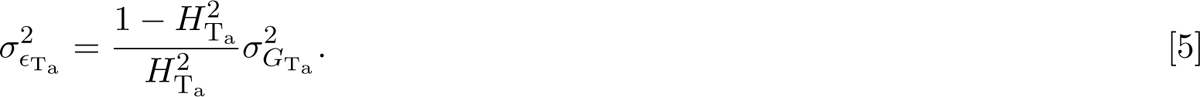

In this study, the trial environments across locations and breeding cycles were assumed to be homogeneous, and therefore *σ*^2^_trialTa_ and *σ*^2^_trialTt_ were fixed. To do so, *σ*^2^_trialTa_ and *σ*^2^_trialTt_ were estimated at SL of C_0_ and were then both assumed fixed for the following cycles. In detail, the ratio of variance components was set for T_t_ as follows: *σ*^2^_GTt_: *σ*^2^_GTt × L_: *σ*^2^_trialTt_ = 1: 1: 0.5, and the corresponding heritability (*H*^2^_Tt_) at each breeding stage was calculated as 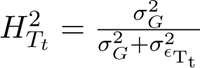. For instance, the *H*^2^ at D clone stage was 0.73. The heritability of *H*^2^_Ta_ was fixed to 0.6. At SL of C_0_, *σ*^2^_GTt_ and *σ*^2^_GTt_ were estimated by the sample variance of TGV_Ta_ and TGV_Tt_, respectively. Then, *σ*^2^_trialTt_ was fixed to ½ of the estimated *σ*^2^_trialTt_. Similarly, *σ*^2^_GTt_ was estimated by equation [5]. However, *σ*^2^_GTt_ and *σ*^2^_GTa_ varied across breeding cycles and *σ*^2^_GTt_ was re-estimated at SL of each cycle. Consequently, *σ*^2^_GTt_ was controlled by the ratio of variance components.

#### 2.4.2 Estimated breeding and genetic values

In this study, a GS model was assumed to be trained for T_t_ on earlier cycles of the breeding program with a prediction accuracy PA. The estimated breeding values for T_t_ obtained from the GS model were estimated by EBV_Tt_ = TBV_Tt_ + *ɛ*_PA_, where TBV_Tt_ were the true breeding values of T_t_, for which only additive effects were considered. *ɛ*_PA_ was the residual value following a normal distribution *N* (0*, σ*^2^), with

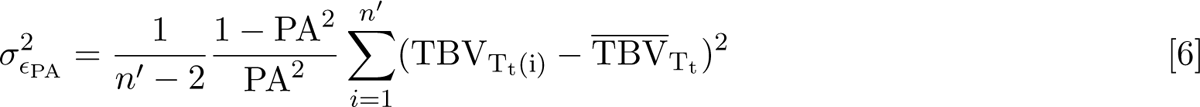

representing the error variance determined by the level of PA, where *n^′^* was the number of genotyped clones, TBV_T (i)_ the TBV_T_ at the *i^th^* genotyped clone, and TBV_Tt_ the average of TBV_Tt_ on all genotyped clones. The estimated genetic values for T_t_ (EGV_Tt_) were obtained by replacing all TBV appearing in this section by TGV.

### 2.5 Economic settings and quantitative genetic parameters

The costs for phenotypic evaluation of T_a_ and T_t_ in one environment were assumed to be 1.4 and 25 e, respectively. The costs for genotypic evaluation per clone were set to 25 e. For the Standard-PS procedure, the total budget in one breeding cycle was 677,500 e. As this strategy served as benchmark, the total budget for all other selection strategies was also fixed to this amount. In this study, we chose the selection strategy GS-SH:A as Optimal-GS (see Wu et al. 2023), and set PA and r to 0.5 and 0.15, respectively, for all selection strategies as well as CS methods. The same number of locations and number of clones at D (N_6_=60) were set as the ones in the Standard-PS (see Wu et al. 2023). The optimal selection proportions achieving the maximum short-term genetic gain and the number of clones at SL for each selection strategy used in this study are summarized in Figure S1.

In order to investigate the interaction effects between different genetic architectures and CS methods on the long-term genetic gain, we considered four different cases of degree of dominance *δ* for T_t_: (1) No dominance effects: *δ*_0_ was set to 0; (2) mild dominance effects: *δ*_1_ was produced across all QTL from *N* (1, 1); (3) moderate dominance effects: *δ*_2_ = 2 *× δ*_1_; (4) strong dominance effects: *δ*_3_ = 3 *× δ*_1_. The true dominance effect at each QTL was then calculated by multiplying the true additive effect by the specific *δ*.

### 2.6 Evaluations

The genetic gain and genetic variability of TGV_Tt_, the genome-wide diversity, as well as the number of QTL where the favorable allele was fixed or lost was estimated and ranked for each scenario in each breeding cycle. The genetic gain was defined as the difference in mean TGV_Tt_ between progenies at D clone stage and the 80 selected candidate parents of C_0_. The level of variability was evaluated by the genetic variance of T_t_, and the level of genomic diversity by the expected heterozygosity (He) (see equation [3]) at D clone stage. The number of QTL where the favorable allele was fixed (= all progenies carrying genotype with AAAA) or lost (= all progenies carrying genotype with aaaa) was calculated among the progenies at D clone stage. To avoid effects due to sampling, all results in this study were based on 30 independent simulation runs.

The efficiency of converting genetic diversity into genetic gain was measured by regressing the realized genetic gain (*y*) on the loss of genetic diversity (*x*), i.e. *y* = *a* + *bx* + *e*, in which the slope (*b*) was efficiency (Gorjanc et al., 2018). In this study, large fluctuations in genetic variance were noticed especially with increased dominance effects. Thus, the realized genetic gain (*y*) was calculated by the difference in averaged genetic gain among 30 simulation runs between C_0_ and C_30_. Similarly, the loss of genetic diversity was computed as the difference in the averaged genetic standard deviation among 30 simulation runs between C_0_ and C_30_.

To assess the accuracy in predicting progeny mean using different mean-based CS methods, we calculated the real progeny mean as the average of all simulated SL progenies at C_0_ and C_30_, respectively. The prediction accuracy was estimated as the correlation between real and predicted progeny mean on an average across 30 simulation runs.

## 3 RESULTS

The mean genetic gain and genetic variance of T_t_, the efficiency of converting genetic diversity into genetic gain, the genome-wide diversity, as well as the number of QTL where the favorable allele was fixed or lost in a long-term tetraploid potato breeding program were assessed considering the following parameters and their interactions: (1) different selection strategies, (2) different CS methods, and (3) different genetic architectures of T_t_, i.e., different degree of dominance.

Regardless of the genetic architectures of T_t_ and under the use of the MPV method, any selection strategy based on the optimal allocation of resources (Optimal-GS and Optimal-PS) had a higher genetic gain than the Standard-PS in both short- and long-term breeding programs (Figure 2a). Furthermore, Optimal-GS was superior to Optimal-PS. An increase in the cycle numbers strengthened this tendency.

Regardless of the selection strategies, CS methods, and genetic architectures of T_t_, an improved genetic gain was observed with increased number of completed breeding cycles (Figures 2a and 5a). However, the additional genetic gain per cycle became smaller at late breeding cycles compared to early ones. This trend as well as the difference in ranking among all assessed CS methods were affected by several parameters: the degree of dominance and weights (*w*_1_ and *w*_2_) of the modified UC. Their details are explained below.

### 3.1 Comparison of CS methods that only consider progeny mean

First, we observed the effects of the implementation of GS on genetic gain using different CS methods only focusing on the progeny mean. In general, any progeny mean predicted by in silico progenies (MEBV-O and MEGV-O) outperformed those predicted by mid-parental performance (MPV, MEBV-P, and MEGV-P) (Figure 2a). Furthermore, the MEGV-O method was superior to the MEBV-O method. The difference between these two CS methods was more obvious with increasing number of breeding cycles as well as an increased the degree of dominance. The latter had stronger influences on genetic gain compared to the former. In addition, the MPV (Optimal-GS) had the highest long-term genetic gain among CS methods based on mid-parental performance. Interestingly, a higher prediction accuracy in predicting progeny mean was observed for the methods based on in silico progenies compared to those based on mid-parental performance (Figure 3).

**Figure 3:**
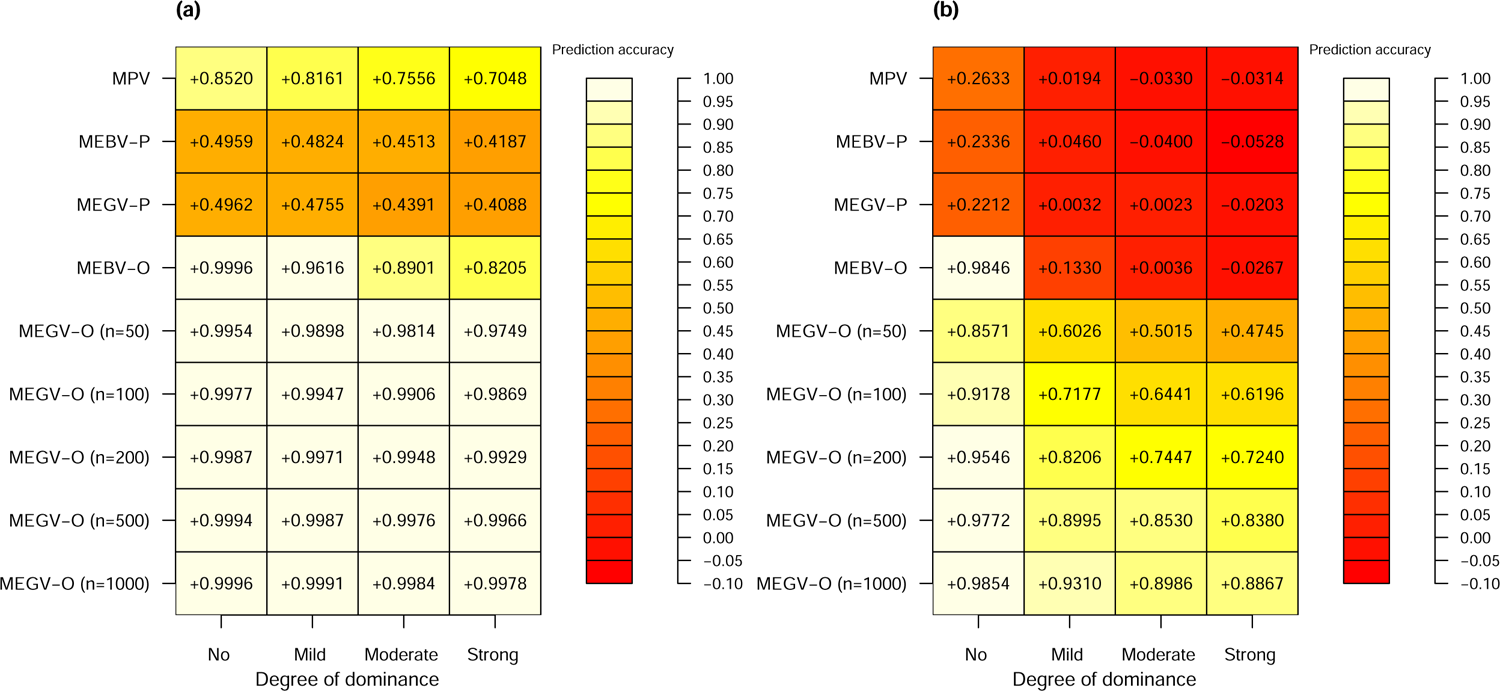
Accuracy to predict progeny mean using the different mean-based cross-selection methods (MPV, MEBV-P, MEGV-P, MEBV-O, and MEGV-O) under different genetic architectures of the target trait. The accuracy was calculated as the correlation between predicted progeny mean and real progeny mean at seedling stage of C_0_ (a) and C_30_ (b), respectively on an average across 30 simulation runs. To examine whether the population size of simulated progenies affects the prediction accuracy using MEGV-O, five different population sizes of the simulated progeny (n = 50, 100, 200, 500, and 1000) were considered.

In contrast to the genetic gain, the genetic variance of T_t_ decreased as the number of breeding cycles increased (Figure 2b). This tendency increased with the reduction of the degree of dominance. Furthermore, the effects of the selection strategies and the CS methods were opposite to the genetic variance in comparison to the genetic gain. As the degree of dominance increased, larger differences and fluctuations in genetic variance among these CS methods and across cycles were observed.

On the other hand, all mean-based CS methods had similar efficiency of converting genetic diversity into genetic gain under the cases without and with low dominance effects (Figure 2c). With increasing dominance effects, MEGV-O did not reach the largest efficiency among all mean-based CS methods. However, its genetic gain was about 1.3 times higher than the CS method achieving the highest efficiency under Optimal-PS.

With increasing number of completed breeding cycles, the genome-wide diversity measured as He decreased (Figure 4a). Simultaneously, the number of QTL where the favorable allele was fixed or lost increased as well (Figure 4b and c). However, a higher degree of dominance reduced this tendency. With an increase of the importance of dominance effects, the CS methods considering additive and dominance effects (MPV, MEGV-P, and MEGV-O) kept a higher He and resulted in a lower number of fixed QTL than those based solely on additive effects (MEBV-P and MEBV-O), especially at late cycles. Furthermore, the MEGV-O method kept the highest He and had the lowest number of fixed QTL among the progeny mean-based CS methods, even though it had the lowest genetic variance and the highest genetic gain. Therefore, MEGV-O was used hereafter as the measurement for the prediction of progeny mean in the weighted methods, i.e., EUC and EUCD.

**Figure 4:**
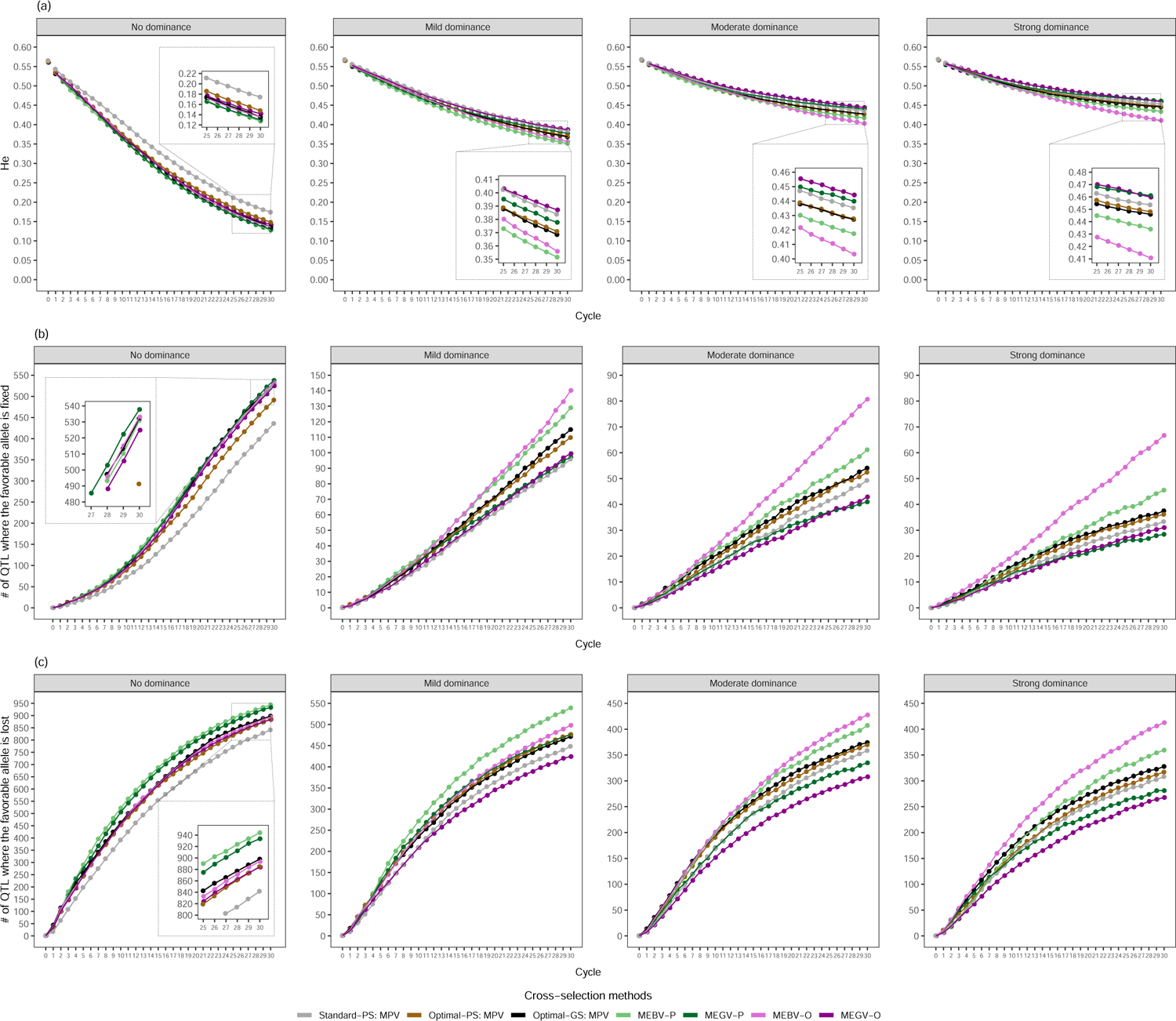
The evolution of genome-wide diversity measured by expected heterozygosity (He) (a), number of QTL where the favorable allele is fixed (b) and lost (c), along the 30 breeding cycles on average across 30 simulation runs. The three parameters were assessed at D clone stage for different selection strategies (Standard-PS, Optimal-PS, and Optimal-GS), different mean-based cross-selection methods (MPV, MEBV-P, MEGV-P, MEBV-O, and MEGV-O), and different genetic architectures of the target trait (no, mild, moderate, and strong dominance effects).

### 3.2 Comparison of CS methods with weights on progeny variance or **genome-wide diversity**

Regardless of the genetic architecture of T_t_, a small or no difference in genetic gain was observed at early cycles among the following CS methods: MEGV-O, EUC, and EUCD with low weights (Figure 5a). As the cycle number increased, the difference became more pronounced. On average across the four levels of dominance effects, EUC_(1,0)_ (=UC) had the highest genetic gain among all EUC approaches (731.01 at C_30_), and was superior to CS methods based only on progeny mean (MEGV-O and MPV methods) (Figure 5a and Table S1). Furthermore, EUCD with a low weight (*w*_2_=50 or 500) yielded the highest genetic gain (734.38 at C_30_).

**Figure 5:**
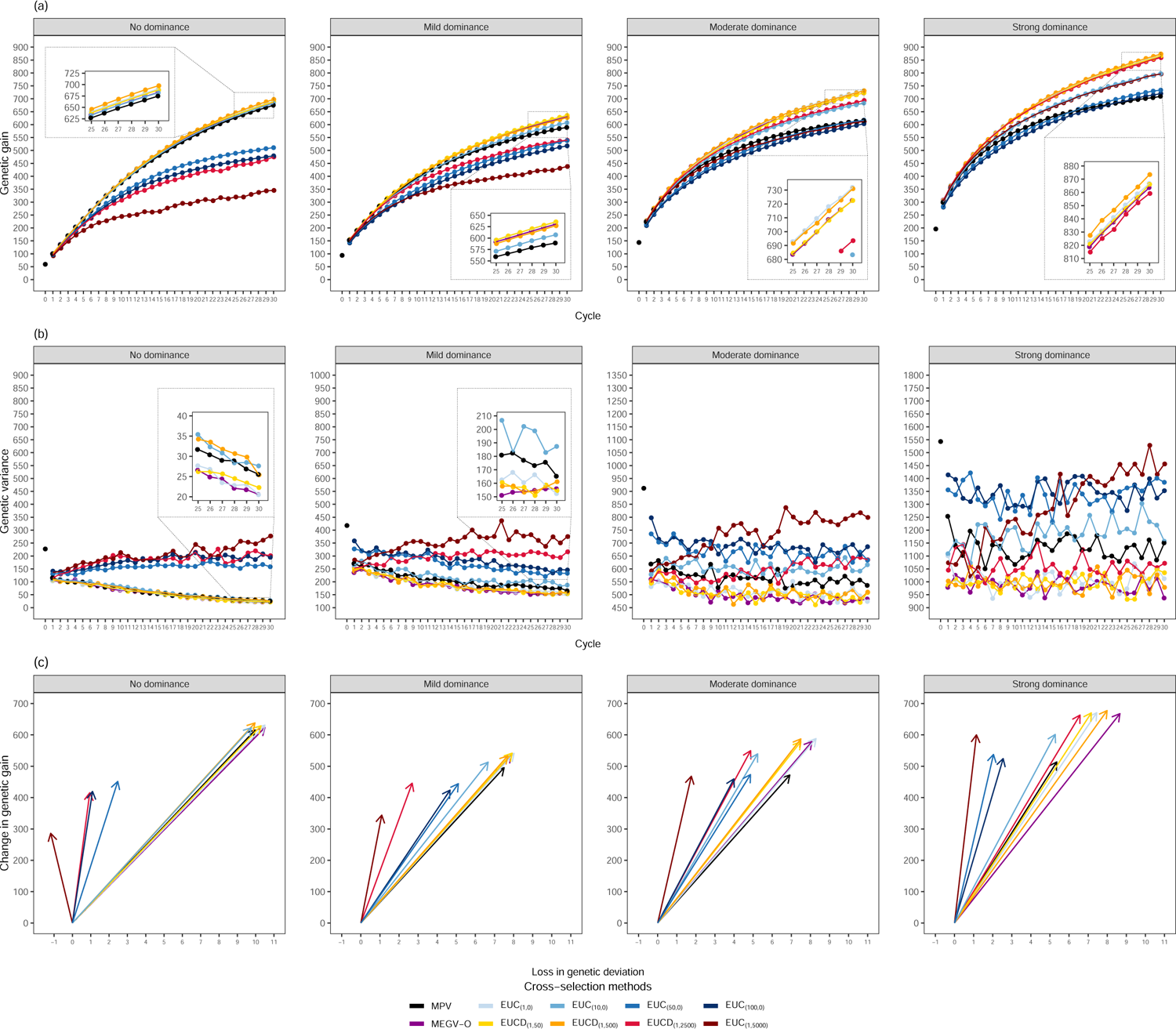
The evolution of genetic gain (a) and genetic variance (b) for the target trait along the 30 breeding cycles on average across 30 simulation runs. The efficiency of converting genetic diversity into genetic gain (c) by regressing the change of genetic gain on the loss of genetic standard deviation between cycle 0 and cycle 30. The three parameters were assessed at D clone stage based on the Optimal-GS selection strategy for different cross-selection methods modified by usefulness criteria (EUC and EUCD), and different genetic architectures of the target trait (no, mild, moderate, and strong dominance effects). The details of EUC and EUCD are shown in Table 1.

We compared four different levels of importance for the variability aspect (being genetic variance or He) in EUC/EUCD on the long term gain of selection. These were called Scale A, B, C, and D (Table 1). Regardless of the genetic architecture of T_t_, no significant difference between the genetic gain of EUCD and EUC was observed when the lowest weights for *w*_1_ and *w*_2_ were considered (i.e. scale A, Figure 5a and Table S1). Furthermore, EUCD_(1,500)_ always outperformed EUC_(10,0)_ (Scale B). With high dominance effects, the EUCDs were superior to the EUCs with high weights, i.e., under Scale C and D.

The ranking and the difference in genetic gain among the above mentioned CS methods were influenced by the degree of dominance (Table S1). EUC and EUCD with high weights ranked better when the importance of dominance effects increased. This was especially true for EUCD. For instance, EUCD_(1,5000)_ had the worst performance under no or mild dominance effects. However, with strong dominance effects, it ranked 7th and outperformed EUC_(50&100,0)_, as well as MPV. While a slow improvement of genetic gain using EUCD_(1,2500)_ was observed under the case without dominance effects, it ranked 5th under the cases with moderate and strong dominance effects. Furthermore, the difference between this CS method and the best one decreased, especially in the case of strong dominance effects.

EUC and EUCD with low weights resulted in high genetic gain but were accompanied by low genetic variance (Figure 5a & b and Table S1). This trend was similar to the mean-based CS methods described in the previous section. In addition, with an increase in cycle numbers, the reduction of genetic variance slowed down, especially for the scenario with strong dominance effects. Differently, high-weighted EUC and EUCD kept relatively high genetic variance and even increased it as the cycle number increased.

The CS methods were also compared with regard to their effects on genetic variance within each scale (Table 1). EUCD resulted also in a higher genetic variance than EUC under Scale C and D, except for the case with strong dominance effects under Scale C. Nevertheless, EUCD resulted in a higher genetic gain than EUC. Furthermore, with strong dominance effects, EUCD_(1,5000)_ kept the highest genetic variance. However, it still performed similarly to EUC_(10,0)_ regarding genetic gain and even had a much higher genetic gain than MPV and EUC_(50&100,0)_.

In general, any EUC and EUCD had a higher efficiency of converting genetic diversity into genetic gain than MEGV-O, especially with increasing importance of dominance effects (Figure 5c). Higher weights for EUC and especially EUCD had a higher efficiency but lower genetic gain compared to lower weights. Differently, as the importance of dominance effects increased, the difference in genetic gain gradually diminished between using CS methods with higher weights (still reaching a higher efficiency) and CS methods with lower weights.

On the other side, along increasing cycles, EUC dramatically decreased He and increased the number of QTL where favorable allele was fixed or lost (Figure 6 and Table S1) and, thus, had similar trends as the mean-based CS methods. These trends were not mitigated a lot as *w*_1_ increased, except for the scenarios with low or no dominance effects. In contrast to EUC, using EUCD obviously slowed down the decline of He, and simultaneously reduced the number of fixed QTL. A greater *w*_2_ increased this tendency.

**Figure 6:**
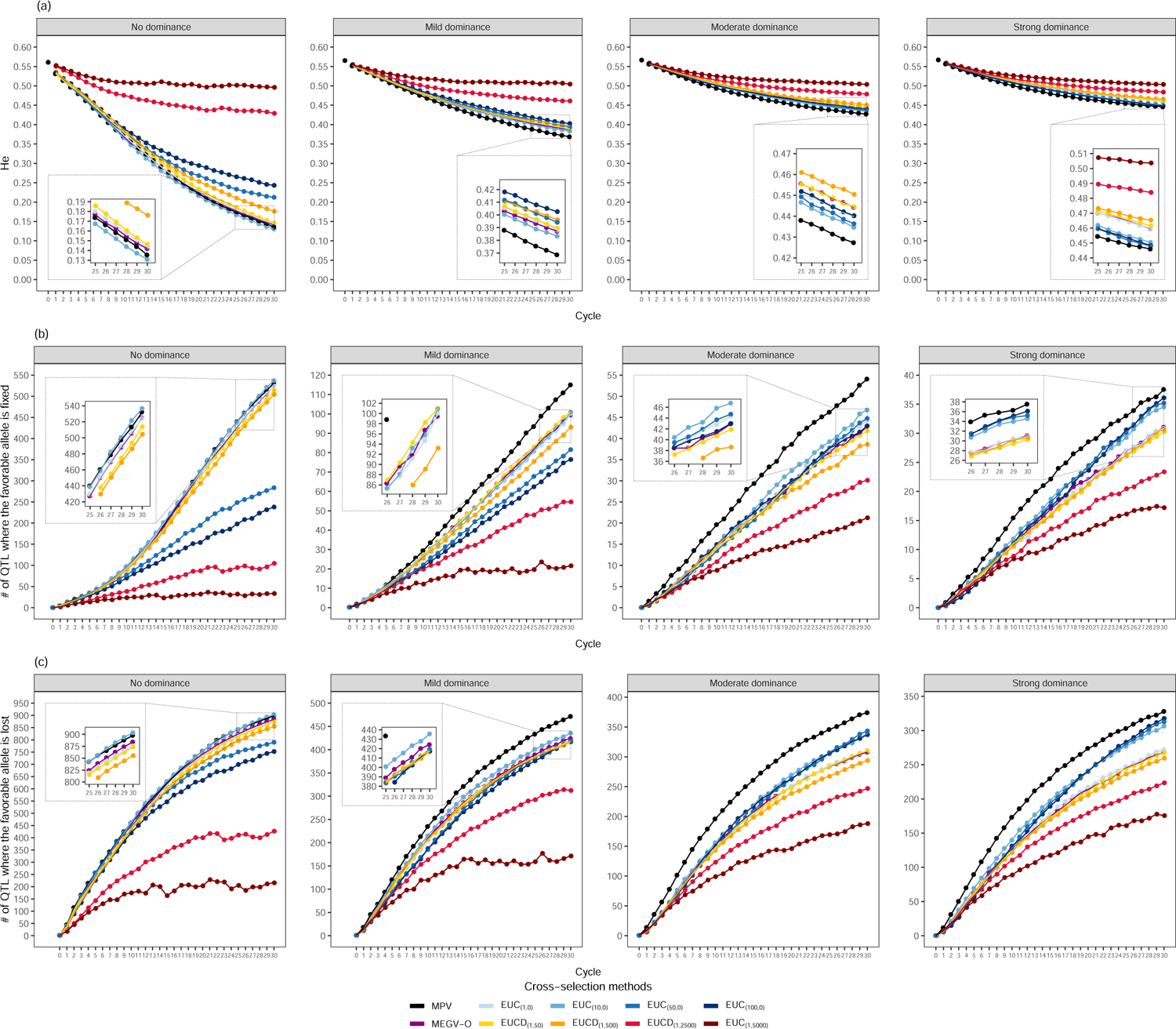
The evolution of genome-wide diversity measured by expected heterozygosity (He) (a), number of QTL where the favorable allele is fixed (b) and lost (c), along the 30 breeding cycles on average across 30 simulation runs. The three parameters were assessed at D clone stage based on the Optimal-GS selection strategy for different cross-selection methods modified by usefulness criteria (EUC and EUCD), and different genetic architectures of the target trait (no, mild, moderate, and strong dominance effects). The details of EUC and EUCD are shown in Table 1.

EUCD kept a higher He and a lower number of fixed QTL than EUC for each scale (Table 1). Furthermore, EUCD with a low *w*_2_ reached a higher He and resulted in a lower number of fixed QTL compared to EUC with a high *w*_1_, especially when dominance effects had a high importance. Across all genetic architectures of T_t_, EUCD with a low *w*_2_ (50 or 500) achieved a high genetic gain and still kept a higher He and a lower number of fixed QTL than the UC and the MEGV-O method. Meanwhile, EUCD’s level of genetic variance remained average. Under strong dominance effects, the genetic gain realized by EUCD with a high *w*_2_ (e.g. EUCD_(1,2500)_) had no significant difference with the highest one reached by EUCD_(1,500)_. However, higher He, genetic variance, as well as efficiency of converting genetic diversity into genetic gain and a lower number of fixed QTL were achieved by EUCD_(1,2500)_ compared to the ones realized by EUCD_(1,500)_.

## 4 DISCUSSIO

Different CS methods accounting or not for diversity have been evaluated in diploid crops to enhance genetic gain (Gaynor et al., 2017; Allier et al., 2019a; Werner et al., 2023). However, the effects of implementing GS and different CS methods in a long-term breeding program for autotetraploid crops with a highly heterozygous genome are lacking. Because of their difference in quantitative genetics compared to diploid inbred or hybrid system, one could expect different outcomes in such analyses. Therefore, we evaluated the efficiency of different CS methods in long-term breeding programs under different genetic architectures via a simulation study.

### 4.1 The effects of different selection strategies on long-term potato **breeding programs**

In this study, we extended an analysis considering the implementation of GS in one breeding cycle (Wu et al., 2023), to study its impact on the long-term genetic gain. Regardless of the genetic architectures and based on MPV as CS method, a higher genetic gain (Figure 2a) was observed in long-term breeding programs with Optimal-PS compared to the benchmark Standard-PS. This follows the trend observed in the study on short-term genetic gain (Wu et al., 2023). The reason is that Optimal-PS had lower selection proportions at B and C clone stages (i.e., higher selection intensities, Figure S1) which were fully based on P_Tt_ selection in comparison with the benchmark procedure. This in turn leads to a higher genetic gain according to the breeder’s equation (Falconer & Mackay, 1996). Furthermore, the selection strategy incorporating GS reached a higher genetic gain than PS did, which can be expected because the former has a higher indirect selection response than the latter at the early stages (Wu et al., 2023). Thus, we compared in the following the performance of the evaluated CS methods using the selection strategy GS-SH:A, i.e., GS was applied at SH and A.

### 4.2 The accuracy of predicting progeny mean

Among the examined mean-based CS methods, the ranking with respect to the maximum genetic gain was MEGV-O *>* MEBV-O *>* MPV *>* MEGV-P and MEBV-P (Figure 2a). This trend was even more pronounced as both the number of completed breeding cycles and the importance of dominance effects increased. One reason might be that the CS methods which rely on simulated offspring can more precisely predict progeny mean compared to mid-parental performance incorporating GS (MEBV-P and MEGV-P) because the former allows to estimate the allele effects more precisely across the progenies of a cross compared to deriving it from parental information. The accuracy of predicting progeny mean (Figure 3) was in complete agreement with our finding about the ranking of the CS methods with respect to their genetic gain. In addition, to examine whether the population size of the simulated progenies affects the degree of prediction accuracy, we varied the number of simulated progeny (n = 50, 100, 200, 500, and 1000). The prediction accuracies among different population sizes of simulated progeny did only vary marginally compared to the ones among different CS methods. Thus, using the CS methods that rely on an in silico cross of simulated offspring results in a higher improvement of genetic gain compared to all CS methods based on mid-parental values (Figure 2a).

Outbred crops have a highly heterozygous genome, which is accompanied by the importance of dominance effects for quantitative traits. However, the proportion of dominance variance components in total genetic variance (including additive and dominance effects) varies depending on the assessed traits and breeding materials. For instance, Endelman et al. (2018) showed that in tetraploid potato dominance variance accounted for 9.4 %, 13.3 %, and 16.4 % of the total genetic variance for the traits specific gravity, yield, and fry color, respectively. In contrast, Thelen et al. (in preparation) showed that dominance variance explained between 0 to 81.1 % of the genetic variance for various agronomic traits. For example, they reported a dominance variance of 50 % for tuber yield, which is considerably higher than the one reported by Endelman et al. (2018). On the other side, the dominance effects in heterozygous species can be partially transmitted from parents to progenies (Gallais, 2003; Endelman et al., 2018; Wolfe et al., 2021; Werner et al., 2023). Therefore, taking into account dominance effects to predict progeny mean can lead to more accurate estimates compared to additive effects only. This was clearly observed in our results based on tetraploid potato: MEGV-O had higher accuracy in predicting progeny mean compared to MEBV-O, especially as the importance of dominance effects increased (Figure 3). It also provided a higher long-term genetic gain, which is in accordance with a former study (Werner et al., 2023). These authors showed that the genetic gain increased when considering both additive and dominance effects to predict cross performance using a formula in a diploid crop. However, our previous statement about the superiority of methods incorporating dominance effects to predict progeny mean was in discordance with our observation that MEGV-P’s genetic gain did not outperform MEBV-P’s genetic gain, despite the fact that only MEGV-P considered dominance effects. One explanation can be that using MEGV-P based on parental dominance effects to capture dominance effects for progenies is an incorrect assumption, leading to a low accuracy in predicting progeny mean, especially with increasing dominance effects (Figure 3).

One surprising aspect was that MPV had the highest genetic gain among all CS methods that relied on mid-parental performance. This observation that phenotypic records outperformed estimated values from a GS model was unexpected, as well as in discordance with former studies in maize breeding programs (Allier et al., 2019a; Sanchez et al., 2023), where MEBV-P reached a higher genetic gain than MPV. One explanation of the superiority of MPV compared to MEBV-P and MEGV-P in our study is that the heritability across the four environments (0.73 at D clone stage of C_0_) used in the first method was higher than the assigned PA (0.5) used in the latter ones. Therefore, according to the breeder’s equation, the MPV can increase more the genetic gain than other CS methods based on mid-parental performance incorporating a GS model. This result was also confirmed by the observed higher accuracy in predicting progeny mean using MPV compared to MEBV-P and MEGV-P (Figure 3).

### 4.3 The limitation of mean-based CS methods

Besides genetic gain, the evaluation of genetic variability across cycles is essential because low genetic variations in breeding materials could limit the genetic gain in the long-term (Falconer & Mackay, 1996). As expected, both the genetic variance of T_t_ and He decreased with increasing cycle numbers (Figures 2b and 4a). At the same time, the number of QTL where the favorable allele was fixed or lost increased as well (Figure 4). The reduction in genetic variance was more pronounced especially for the CS methods reaching the higher genetic gain. The high accuracy in predicting progeny mean that leads to the quick accumulation of favorable alleles (Figure 4b) can be one reason for this observation. Moreover, the Bulmer effect (Bulmer, 1971), which reduces the proportion of genetic variance due to linkage disequilibrium between trait-coding polymorphisms (Grevenhof et al., 2012), can further explain this result. In order to assess the potential importance of the Bulmer effect, we calculated the maximum genetic gain as the difference between the maximum genetic value and mean genetic values among the 80 selected candidate parents of C_0_, where the maximum genetic value was obtained by summing up the maximum genetic values among the five genotypes of each QTL (Table 2) across the 2,000 QTL. The genetic gain of the mean-based CS methods gradually closed up to the maximum genetic gain under the case without dominance effects (Figure S2), implying that the influence of the Bulmer effect was not high.

Overall, focusing on mean performance only to select new crosses could lead to a plateau for the genetic gain with increasing cycle numbers. Therefore, CS methods considering the maintenance of diversity while maximizing long-term genetic gain are required.

### 4.4 The efficiency of CS methods to balance genetic gain and **maintenance of diversity**

Besides paying attention to high progeny mean, a high variance in progenies is of fundamental importance for the response to selection. The UC of a cross considers these aspects and has been used to predict the mean performance of the upper fraction of its progeny, considering the genetic variance, the heritability as well as the selection intensity (Allier et al., 2019a; Sanchez et al., 2023). Thus, this method could improve the genetic gain compared to mean-based CS methods, which is confirmed in our study (Figure 5a and Table S1). However, while we observed a slightly higher genetic gain using the UC compared to the MEGV-O method (Figure 5 and Table S1), the genetic variance or He were the same for UC and MEGV-O method.

Furthermore, the difference in the genetic gain between UC and MEGV-O was not statistically significant, which is contradictory to the results of former studies in diploid crops (Lehermeier et al., 2017; Sanchez et al., 2023). This could be explained by the lower PA (0.5) and selection intensity (1.75) used in the present study, compared to a high heritability (1) and selection intensity (2.06) in Sanchez et al. (2023). Lehermeier et al. (2017) also showed that higher heritability and selection intensity lead to a higher advantage of the UC versus other methods.

On the other hand, the variance in the progeny mean was much higher (*∼* 90 times) than the variance in the progeny standard deviation in our study. This is in accordance with former studies (Zhong & Jannink, 2007; Lado et al., 2017), leading to no difference between UC and progeny mean. Thus, one way to strengthen the importance of the genetic variance in the progeny could be to increase the weight of the genetic variance or to add an extra variation measurement to the UC.

The genetic diversity of a cross can be quantified by the genetic variance of a trait, but also on a genome-wide scale by the expected heterozygosity (He) estimated from molecular genetic information. Therefore, in addition to the weight on genetic variance of T_t_, i.e., EUC, one could consider weighting He to integrate another level of diversity to balance genetic gain. This is because the latter considers the level of whole genomic variation instead of being restricted to the variation of specific loci linked to QTL of T_t_ like the former. In our study, on average across the four different genetic architectures (from no to strong dominance), EUCD_(1,50|500)_ reached the maximum genetic gain among all assessed EUCDs and a slightly higher long-term genetic gain compared to UC (Figure 5a and Table S1). Meanwhile, EUCD_(1,50|500)_ kept a certain degree of genetic variance, a slightly higher He, as well as a bit lower number of fixed QTL compared to UC (Figures 5 and 6). This confirmed our expectation, as EUCD keeps the advantage of the UC and preserves a certain genome-wide diversity by accounting for He simultaneously, which in turn helps to efficiently convert genetic variability into long-term genetic gain (Figure 5c).

While EUCD with a high weight kept a higher genetic variance and He along the cycles as well as had a lower number of fixed QTL, it was accompanied by a reduction of the long-term genetic gain compared to EUCD with a low weight. This was not surprising because a high weight on diversity means to minimize the loss of diversity after selection. Allier et al. (2019a) had a similar approach accounting for different weights of penalty on He to balance between maximal genetic gain and minimal loss of diversity during the selection of new crosses. Their result also indicated that a stronger penalty on diversity reduced the improvement of genetic gain but kept higher diversity. However, this trend of lower genetic gain with a higher *w*_2_ gradually diminished as the degree of dominance increased in our study, implying different weights should be fitted to different genetic architectures when using EUCD, as different degrees of dominance appear in the agronomic traits of potato in experimental studies.

Although our proposed method EUCD cannot manage to reach a significant improvement in genetic gain compared to EUC and MEGV-O, it keeps a higher genome-wide diversity, which can balance maximal genetic gain and minimal loss of diversity in the process of selecting new crosses. Preserving diversity is very important in long-term breeding programs because it provides opportunities for breeders to promptly adjust the goals of the breeding programs in response to new requests such as changes in climate and human usage and to develop new varieties adapted to biotic and abiotic stresses. Therefore, for the improvement of the long-term breeding program, potato breeders should choose a proper weight on He accounting to their parameters for a subsequent long-term improvement in genetic gain and nevertheless adaptability of the breeding program. In detail, to reach a high long-term genetic gain but simultaneously keep a certain diversity, EUCD_(1,50|500)_ can be used for cases with no, mild, and moderate dominance effects, where EUCD_(1,2500)_ seems to be appropriate for cases with strong dominance effects. However, EUCD_(1,2500)_ or EUCD_(1,5000)_ can be utilized if the main breeding goals are to keep maximum diversity and to reach a certain genetic gain for the cases with moderate or strong dominance effects. Therefore, the choice of the most appropriate weight on diversity in EUCD depends not only on the genetic architecture of T_t_, but also on the breeder’s objectives.

### 4.5 Assumptions of the present study

In this study, we assume that the parental haplotype phase is known, and therefore, the progeny variance can be predicted by in silico progenies (Bernardo, 2014; Mohammadi et al., 2015; Miller et al., 2023). However, also with current methodology (e.g., Sun et al. 2022), the assessment of the haplotype phase is costly. Thus, in current breeding programs, the possibility of estimating the progeny mean is based on mid-parent performance. In this study, MPV had a higher accuracy in predicting progeny mean compared to MEGV-P or MEBV-P because the heritability (0.73) is higher than the PA (0.5). However, if heritability is lower than PA, the advantage of MPV compared to MEBV-P and MEGV-P will disappear. For example, the heritability at early breeding stages is lower than the one at late breeding stages, because the former has less experimental locations and replications than the latter. Therefore, if the candidate parents are selected from early breeding stages, the superiority of MPV over MEBV-P or MEGV-P will diminish.

Wolfe et al. (2021) and Werner et al. (2023) predicted the progeny mean with the formula based on allele frequencies of parents and considering additive and dominance effects from Falconer & Mackay (1996) in heterozygous diploid crops. Although Wolfe et al. (2021) showed no improvement in prediction accuracy of the progeny mean using MEGV estimated by the formula compared to MEBV estimated by mid-parental values in an experimental dataset, the results of the simulation study of Werner et al. (2023) indicated that the genetic gain was improved using MEGV estimated by the formula to select crosses especially for traits with dominance effects. Therefore, one possibility to improve the prediction of progeny mean in future research is to develop the formula to estimate progeny mean and variance in autotetraploid species. Furthermore, He*_per−cross_* based on simulated progenies is highly correlated with He*_per−cross_* based on parental genotypic information (data not shown). Thus, the lack of information about haplotype phase does not affect the ability to quantify genome-wide diversity of a cross.

An alternative method to consider genome-wide diversity while selecting new crosses for the next breeding cycle was developed by Gorjanc et al. (2018) and Allier et al. (2019a). Their approach is called OCS and is based on an optimization algorithm. This approach provided a high genetic gain as well as kept a high diversity. However, this method required the optimization process to search for an optimal group of crosses, leading to extremely intensive computational calculations compared to our proposed EUCD methods. This difference in computational time requirement is even more pronounced with an increasing number of markers, repetitions, and candidates. Our study considered between 2 to 24 times more SNP and triple repetition numbers compared to the former studies. In addition, the number of all possible solutions to select 300 crosses from 1790 possible cross combinations in this study is infinite, which was far greater than the ones in former studies. Thus, OCS has not been assessed yet in this study. However, the comparison of performance between the two methods requires further research.

### 4.6 Summary

The present study demonstrated that implementing GS with optimal selection intensity per stage enhances both short- and long-term gain from selection compared to a typical tetraploid potato breeding program solely based on PS. In addition, for autotetraploid and heterozygous crops, the prediction of progeny mean considering not only additive but also dominance effects (MEGV-O) is advantageous. This approach results in the highest prediction accuracy to predict progeny mean and have the highest genetic gain among all mean-based CS methods. Furthermore, combining UC and genome-wide diversity (EUCD) by a linear combination reached the same level of long-term genetic gain in a tetraploid potato breeding program. However, it simultaneously preserved a higher diversity, a certain degree of genetic variance, as well as a lower number of fixed QTL compared to MEGV-O and UC. In our results, although EUCD with a low weight can reach the highest genetic gain, different genetic architectures of T_t_ and the breeder’s objectives require choosing different weights on genome-wide diversity to achieve a high genetic gain and simultaneously preserve sufficient diversity. These results can provide breeders with a concrete method to improve their potato breeding programs.

## List of abbreviations

Abbreviations: Full name

PS: phenotypic selection

GS: genomic selection

*µ*: progeny mean

*i*: selection intensity

*H*: square root of the heritability

*σ_G_*: square root of the progeny variance

He: expected heterozygosity

X: cross stage

SL: seedling stage

SH: single hills stage

B: A clone stage

C: B clone stage

D: C clone stage

E: D clone stage

C_0_; C_1_; C*_t_*: burn-in cycle, that is cycle 0; cycle 1; cycle t

QTL: quantitative trait loci

Standard-PS: a scheme following a standard potato breeding program relying exclusively on PS

Optimal-PS: a scheme relying on PS but where the optimal selection intensities during the selection process were determined to maximize genetic gain

Optimal-GS: a scheme based on both PS and GS where the optimal selection intensities during the selection process were determined to maximize genetic gain

p*_i_*: selection proportion at the *i^th^*stage

T_a_: auxiliary trait

T_t_: target trait

TBV: true breeding values (additive effects)

TGV: true genetic values (additive and dominance effects)

EBV: estimated breeding values (additive effects)

EGV: estimated genetic values (additive and dominance effects)

P: phenotypic values

PA: prediction accuracy

OCS: optimal cross-selection

CS: method cross-selection method

MPV: mean phenotypic values of the two parents

MEBV-P: mean estimated breeding values of the two parents

MEGV-P: mean estimated genetic values of the two parents

MEBV-O: mean estimated breeding values among simulated offspring

MEGV-O: mean estimated genetic values among simulated offspring

UC: usefulness criterion (Schnell & Utz, 1975)

EUC: extended usefulness criterion, incorporating different weight (*w*_1_) on the progeny variance

EUCD: extended usefulness criterion incorporating genomic diversity index (UC + different weight – *w*_2_ on genome-wide diversity, which is quantified by He)

## ACKNOWLEDGMENTS

Computational infrastructure and support were provided by the Centre for Information and Media Technology (ZIM) at Heinrich Heine University Düsseldorf. This study was funded by the Federal Ministry of Food and Agriculture/Fachagentur Nachwachsende Rohstoffe (grantID 22011818 – PotatoTools and grantID 2222NR078A – PotatoPredict). The funders had no influence on study design, the collection, analysis and interpretation of data, the writing of the manuscript, and the decision to submit the manuscript for publication.

## SUPPORTING INFORMATION

Additional supporting information can be found online in the Supporting Information section at the end of this article.

## DATA AVAILABILITY

The datasets generated or analyzed during this study and R scripts are available from the authors upon request.

## AUTHOR CONTRIBUTIONS

BS and DVI designed and coordinated the study; PYW performed the analyses; SH, KM, and VP provided details about breeding schemes; PYW, BS, and DVI wrote the manuscript. All authors read and approved the final manuscript.

## SUPPLEMENTARY MATERIAL

**Table S1:** The mean and standard deviation (sd) of the genetic gain and the genetic variance for the target trait, as well as genome-wide diversity measured by expected heterozygosity (He) at cycle 30 across 30 simulation runs. Simulations were based on the Optimal-GS selection strategy for different cross-selection methods (MPV, MEGV-O, EUC and EUCD), and different genetic architectures of the target trait (no, mild, moderate, and strong dominance effects). The details of EUC and EUCD are shown in Table 1.

| Case | Scale | Cross-selection method | Genetic gain |  |  |  | Genetic variance |  |  | He |  |  |
| --- | --- | --- | --- | --- | --- | --- | --- | --- | --- | --- | --- | --- |
|  |  |  | mean | sd | rank<sup>1</sup> | group<sup>2</sup> | mean | sd | rank<sup>1</sup> | mean | sd | rank<sup>1</sup> |
| No dominance | - | Optimal-GS: MPV | 674.86 | 45.38 | 6 | b | 25.54 | 7.17 | 6 | 0.1354 | 0.0236 | 9 |
|  |  | MEGV-O | 682.91 | 36.83 | 4 | ab | 20.55 | 7.12 | 9 | 0.1419 | 0.0211 | 8 |
|  | A | EUC<sub>(1,0)</sub>=UC | 690.42 | 34.25 | 2 | ab | 20.50 | 6.79 | 10 | 0.1438 | 0.0241 | 7 |
|  |  | EUCD<sub>(1,50)</sub> | 688.26 | 37.24 | 3 | ab | 22.27 | 5.59 | 8 | 0.1462 | 0.0241 | 6 |
|  | B | EUC<sub>(10,0)</sub> | 682.03 | 35.41 | 5 | ab | 27.62 | 7.19 | 5 | 0.1307 | 0.0216 | 10 |
|  |  | EUCD<sub>(1,500)</sub> | 697.52 | 35.45 | 1 | a | 25.41 | 5.29 | 7 | 0.1762 | 0.0193 | 5 |
|  | C | EUC<sub>(50,0)</sub> | 510.83 | 41.30 | 7 | c | 158.38 | 37.96 | 4 | 0.2125 | 0.0255 | 4 |
|  |  | EUCD<sub>(1,2500)</sub> | 474.79 | 32.86 | 9 | d | 201.18 | 59.78 | 2 | 0.4291 | 0.0106 | 2 |
|  | D | EUC<sub>(100,0)</sub> | 479.07 | 31.80 | 8 | d | 195.13 | 45.51 | 3 | 0.2432 | 0.0287 | 3 |
|  |  | EUCD<sub>(1,5000)</sub> | 345.34 | 29.90 | 10 | e | 277.26 | 152.84 | 1 | 0.4962 | 0.0107 | 1 |
| Mild dominance | - | Optimal-GS: MPV | 589.13 | 24.70 | 6 | b | 165.21 | 29.62 | 6 | 0.3686 | 0.0146 | 10 |
|  |  | MEGV-O | 630.46 | 21.73 | 3 | a | 155.91 | 32.38 | 8 | 0.3871 | 0.0120 | 8 |
|  | A | EUC<sub>(1,0)</sub>=UC | 634.86 | 20.86 | 2 | a | 152.32 | 29.24 | 10 | 0.3880 | 0.0132 | 7 |
|  |  | EUCD<sub>(1,50)</sub> | 635.53 | 18.70 | 1 | a | 154.53 | 28.70 | 9 | 0.3896 | 0.0107 | 6 |
|  | B | EUC<sub>(10,0)</sub> | 607.07 | 24.09 | 5 | b | 187.34 | 32.32 | 5 | 0.3833 | 0.0130 | 9 |
|  |  | EUCD<sub>(1,500)</sub> | 627.08 | 18.74 | 4 | a | 161.12 | 36.10 | 7 | 0.3963 | 0.0103 | 4 |
|  | C | EUC<sub>(50,0)</sub> | 538.28 | 19.13 | 8 | c | 233.10 | 46.12 | 4 | 0.3942 | 0.0122 | 5 |
|  |  | EUCD<sub>(1,2500)</sub> | 539.83 | 21.64 | 7 | c | 316.13 | 86.45 | 2 | 0.4611 | 0.0075 | 2 |
|  | D | EUC<sub>(100,0)</sub> | 517.75 | 26.58 | 9 | d | 245.84 | 39.13 | 3 | 0.4026 | 0.0147 | 3 |
|  |  | EUCD<sub>(1,5000)</sub> | 437.93 | 21.82 | 10 | e | 375.27 | 101.12 | 1 | 0.5045 | 0.0096 | 1 |
|  |  |  | mean | sd | rank<sup>1</sup> | group<sup>2</sup> | mean | sd | rank<sup>1</sup> | mean | sd | rank<sup>1</sup> |
| Moderate dominance | - | Optimal-GS: MPV | 615.59 | 29.91 | 8 | cd | 536.13 | 110.22 | 6 | 0.4273 | 0.0102 | 10 |
|  |  | MEGV-O | 722.54 | 25.61 | 3 | a | 483.54 | 97.25 | 9 | 0.4442 | 0.0087 | 5 |
|  | A | EUC<sub>(1,0)</sub>=UC | 731.97 | 24.21 | 1 | a | 473.95 | 96.69 | 10 | 0.4400 | 0.0072 | 7 |
|  |  | EUCD<sub>(1,50)</sub> | 722.47 | 26.83 | 4 | a | 508.45 | 98.52 | 8 | 0.4446 | 0.0088 | 4 |
|  | B | EUC<sub>(10,0)</sub> | 683.29 | 21.74 | 6 | b | 616.05 | 115.14 | 5 | 0.4346 | 0.0096 | 9 |
|  |  | EUCD<sub>(1,500)</sub> | 731.00 | 25.33 | 2 | a | 510.10 | 108.94 | 7 | 0.4504 | 0.0086 | 3 |
|  | C | EUC<sub>(50,0)</sub> | 617.29 | 29.37 | 7 | c | 633.84 | 104.73 | 4 | 0.4364 | 0.0110 | 8 |
|  |  | EUCD<sub>(1,2500)</sub> | 693.48 | 28.76 | 5 | b | 635.23 | 120.92 | 3 | 0.4788 | 0.0054 | 2 |
|  | D | EUC<sub>(100,0)</sub> | 603.47 | 27.35 | 10 | d | 685.91 | 188.11 | 2 | 0.4402 | 0.0097 | 6 |
|  |  | EUCD<sub>(1,5000)</sub> | 611.82 | 27.36 | 9 | cd | 799.24 | 118.21 | 1 | 0.5038 | 0.0060 | 1 |
| Strong dominance | - | Optimal-GS: MPV | 709.57 | 29.15 | 10 | d | 1150.66 | 223.20 | 5 | 0.4459 | 0.0089 | 10 |
|  |  | MEGV-O | 863.63 | 37.04 | 4 | a | 937.20 | 192.78 | 10 | 0.4598 | 0.0082 | 6 |
|  | A | EUC<sub>(1,0)</sub>=UC | 866.76 | 34.68 | 2 | a | 1012.18 | 189.86 | 8 | 0.4601 | 0.0074 | 5 |
|  |  | EUCD<sub>(1,50)</sub> | 866.11 | 25.14 | 3 | a | 1035.21 | 240.35 | 7 | 0.4616 | 0.0092 | 4 |
|  | B | EUC<sub>(10,0)</sub> | 797.36 | 33.52 | 6 | b | 1157.18 | 237.24 | 4 | 0.4504 | 0.0065 | 7 |
|  |  | EUCD<sub>(1,500)</sub> | 873.48 | 28.85 | 1 | a | 978.41 | 199.76 | 9 | 0.4654 | 0.0067 | 3 |
|  | C | EUC<sub>(50,0)</sub> | 733.52 | 41.32 | 8 | c | 1385.26 | 240.45 | 2 | 0.4479 | 0.0085 | 9 |
|  |  | EUCD<sub>(1,2500)</sub> | 859.17 | 36.96 | 5 | a | 1071.70 | 230.82 | 6 | 0.4840 | 0.0054 | 2 |
|  | D | EUC<sub>(100,0)</sub> | 720.28 | 36.26 | 9 | cd | 1352.71 | 289.42 | 3 | 0.4484 | 0.0107 | 8 |
|  |  | EUCD<sub>(1,5000)</sub> | 796.02 | 25.44 | 7 | b | 1456.23 | 291.13 | 1 | 0.5037 | 0.0047 | 1 |

<sup>1</sup> The number after the sd represent the rank across these cross-selection methods within a specific genetic architecture.
<sup>2</sup> The letters after the rank represent the significance groups ( $P < 0.05$ ) across these cross-selection methods within a specific genetic architecture.

**Figure S1:**
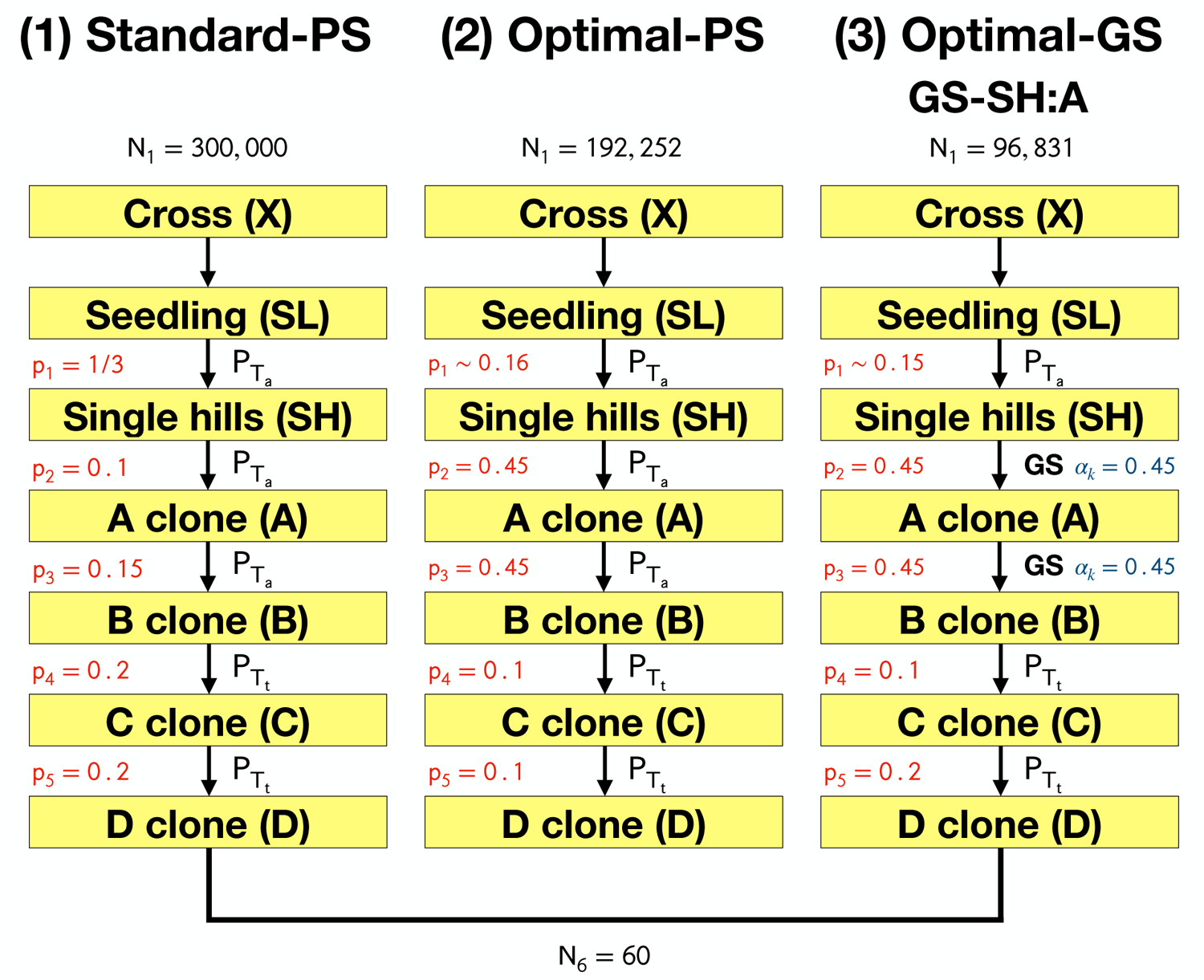
Graphical illustration of the three selection strategies: (1) Standard-PS, where PS is phenotypic selection, (2) Optimal-PS, and (3) Optimal-GS, where genomic selection (GS) is applied at SH and A stages (GS-SH:A) with the prediction accuracy of the GS model of 0.5 and the correlation between the two traits of 0.15. p_1_ to p_5_ are the selected proportions from SL to SH, SH to A, A to B, B to C, and C to D, respectively, where SL, SH, A, B, C, and D represent the stages of seedling, single hills, A, B, C, and D clones. The selected proportions for the strategy (1) Standard-PS follow the standard potato breeding program (Wu et al., 2023). The optimal selected proportions for (2) Optimal-PS and (3) Optimal-GS are determined by achieving the maximum short-term genetic gain. *α_k_* and N_1_ are the weight of genomic selection relative to phenotypic selection and the number of clones at seedling stages, respectively.

**Figure S2:**
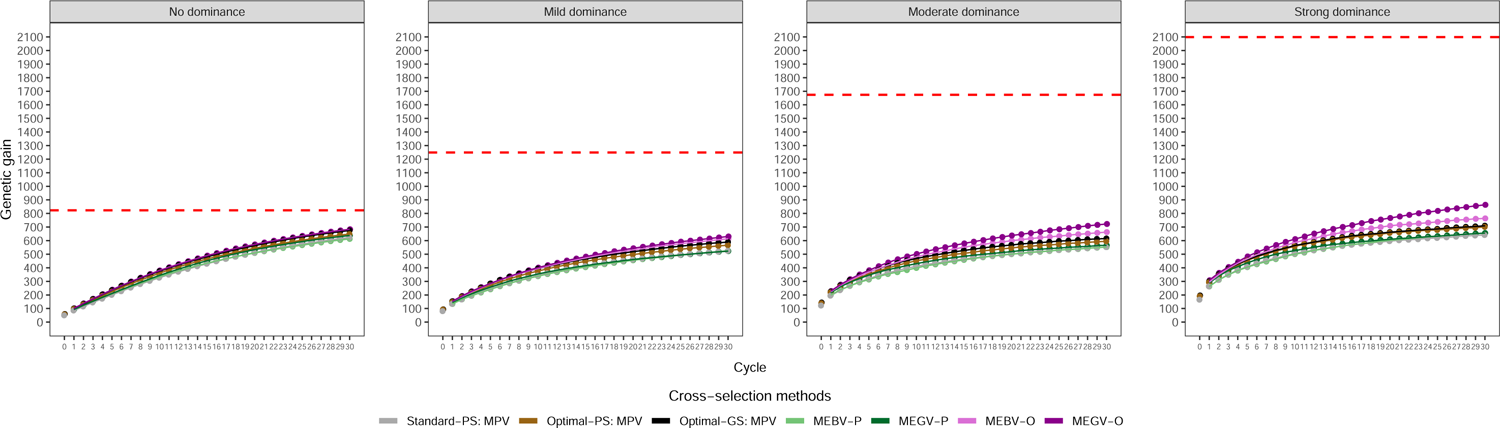
The comparison of maximum genetic gain (see discussion for details – dashed red line) and the genetic gain of different mean-based cross-selection methods across different genetic architectures of the target trait (no, mild, moderate, and strong dominance effects).

## REFERENCES

Abberton M, Batley J, Bentley A, Bryant J, Cai H, Cockram J, Costa de Oliveira A, Cseke L J, Dempewolf H, De Pace C, Edwards D, Gepts P, Greenland A, Hall A E, Henry R, Hori K, Howe G T, Hughes S, Humphreys M, Lightfoot D, Marshall A, Mayes S, Nguyen H T, Ogbonnaya F C, Ortiz R, Paterson A H, Tuberosa R, Valliyodan B, Varshney R K, & Yano M (2016), Global agricultural intensification during climate change: a role for genomics. Plant Biotechnology Journal 14(4):1095–1098

Alemu A, Astrand J, Montesinos-Ĺopez O A, yŚanchez J I, Fernández-Gónzalez J, Tadesse W, Vetukuri R R, Carlsson A S, Ceplitis A, Crossa J, Ortiz R, & Chawade A (2024), Genomic selection in plant breeding: key factors shaping two decades of progress. Molecular Plant 17:552–578

Allier A, Lehermeier C, Charcosset A, Moreau L, & Teyssèdre S (2019a), Improving short-and long-term genetic gain by accounting for within-family variance in optimal cross-selection. Frontiers in Genetics 10:1006

Allier A, Moreau L, Charcosset A, Teyssèdre S, & Lehermeier C (2019b), Usefulness criterion and post-selection parental contributions in multi-parental crosses: application to polygenic trait introgression. G3: Genes—Genomes—Genetics 9(5):1469– 1479

Bernardo R (2014), Genomewide selection of parental inbreds: classes of loci and virtual biparental populations. Crop Science 54(6):2586–2595

Bonk S, Reichelt M, Teuscher F, Segelke D, & Reinsch N (2016), Mendelian sampling covariability of marker effects and genetic values. Genetics Selection Evolution 48(1):1–11

Brown J, & Caligari P D (1989), Cross prediction in a potato breeding programme by evaluation of parental material. Theoretical and Applied Genetics 77(2):246–252

Bulmer M G (1971), The effect of selection on genetic variability. The American Naturalist 105:201–211

Daetwyler H D, Hayden M J, Spangenberg G C, & Hayes B J (2015), Selection on optimal haploid value increases genetic gain and preserves more genetic diversity relative to genomic selection. Genetics 200(4):1341–1348

Endelman J B, Carley C A, Bethke P C, Coombs J J, Clough M E, da Silva W L, Jong W S D, Douches D S, Frederick C M, Haynes K G, Holm D G, Miller J C, Muñoz P R, Navarro F M, Novy R G, Palta J P, Porter G A, Rak K T, Sathuvalli V R, Thompson A L, & Yencho G C (2018), Genetic variance partitioning and genome-wide prediction with allele dosage information in autotetraploid potato. Genetics 209:77–87

Falconer D S, & Mackay T F C (1996), Introduction to quantitative genetics. Longman group, Essex, UK, 4 edition

Fŕona D, Szendeŕak J, & Harangi-Ŕakos M (2019), The challenge of feeding the world. Sustainability (Switzerland) 11(20):5816

Gallais A (2003), Quantitative genetics and breeding methods in autopolyploid plants. Institut national de la recherche agronomique

Gaynor R C, Gorjanc G, Bentley A R, Ober E S, Howell P, Jackson R, Mackay I J, & Hickey J M (2017), A two-part strategy for using genomic selection to develop inbred lines. Crop Science 57(5):2372–2386

Gaynor R C, Gorjanc G, & Hickey J M (2021), AlphaSimR: an R package for breeding program simulations. G3: Genes—Genomes—Genetics 11(2)

Gorjanc G, Gaynor R C, & Hickey J M (2018), Optimal cross selection for longterm genetic gain in two-part programs with rapid recurrent genomic selection. Theoretical and Applied Genetics 131(9):1953–1966

Grevenhof E M V, Arendonk J A V, & Bijma P (2012), Response to genomic selection: the Bulmer effect and the potential of genomic selection when the number of phenotypic records is limiting. Genetics Selection Evolution 44:1–10

Grüneberg W, Mwanga R, Andrade M, & Espinoza J (2009), Selection methods. Part 5: breeding clonally propagated crops. Plant Breeding and Farmer Participation, (edited by S Ceccarelli, E P Guimarães, & E Weltzien), pages 275–322, Food and Agriculture Organization of the United Nations (FAO)

Heffner E L, Lorenz A J, Jannink J L, & Sorrells M E (2010), Plant breeding with genomic selection: gain per unit time and cost. Crop Science 50(5):1681–1690

Jannink J L (2010), Dynamics of long-term genomic selection. Genetics Selection Evolution 42:1–11

Kinghorn B P (2011), An algorithm for efficient constrained mate selection. Genetics Selection Evolution 43(1):1–9

Lado B, Battenfield S, Guzmán C, Quincke M, Singh R P, Dreisigacker S, Peña R J, Fritz A, Silva P, Poland J, & Gutíerrez L (2017), Strategies for selecting crosses using genomic prediction in two wheat breeding programs. The Plant Genome 10

Lehermeier C, Teyssèdre S, & Schön C C (2017), Genetic gain increases by applying the usefulness criterion with improved variance prediction in selection of crosses. Genetics 207(4):1651–1661

Lubanga N, Massawe F, Mayes S, Gorjanc G, & Bančič J (2022), Genomic selection strategies to increase genetic gain in tea breeding programs. The Plant Genome page e20282

Meuwissen T H, Hayes B J, & Goddard M E (2001), Prediction of total genetic value using genome-wide dense marker maps. Genetics 157(4):1819–1829

Miller M J, Song Q, Fallen B, & Li Z (2023), Genomic prediction of optimal cross combinations to accelerate genetic improvement of soybean (Glycine max). Frontiers in Plant Science 14:1–12

Mohammadi M, Tiede T, & Smith K P (2015), PopVar: a Genome-Wide procedure for predicting genetic variance and correlated response in biparental breeding populations. Crop Science 55(5):2068–2077

Muleta K T, Pressoir G, & Morris G P (2019), Optimizing genomic selection for a sorghum breeding program in Haiti: a simulation study. G3: Genes—Genomes—Genetics 9:391–401

Osthushenrich T, Frisch M, & Herzog E (2017), Genomic selection of crossing partners on basis of the expected mean and variance of their derived lines. PLOS ONE 12(12):e0188839

Sanchez D, Sadoun S B, Mary-Huard T, Allier A, Moreau L, & Charcosset A (2023), Improving the use of plant genetic resources to sustain breeding programs’ efficiency. Proceedings of the National Academy of Sciences of the United States of America 120(14):e2205780119

Schnell F, & Utz H (1975), F1 Leistung und Elternwahl in der Zuchtung von Selbst-befruchtern. Ber Arbeitstag Arbeitsgem Saatzuchtleiter. Gumpenstein, Ö sterreich, pages 243–248

Sun H, Jiao W B, Krause K, Campoy J A, Goel M, Folz-Donahue K, Kukat C, Huettel B, & Schneeberger K (2022), Chromosome-scale and haplotype-resolved genome assembly of a tetraploid potato cultivar. Nature Genetics 54:342–348

Werner C R, Gaynor R C, Sargent D J, Lillo A, Gorjanc G, & Hickey J M (2023), Genomic selection strategies for clonally propagated crops. Theoretical and Applied Genetics 136(4):1–17

Wolfe M D, Chan A W, Kulakow P, Rabbi I, & Jannink J L (2021), Genomic mating in outbred species: predicting cross usefulness with additive and total genetic covariance matrices. Genetics 219(3)

Wu P Y, Stich B, Renner J, Muders K, Prigge V, & van Inghelandt D (2023), Optimal implementation of genomic selection in clone breeding programs—Exemplified in potato: I. Effect of selection strategy, implementation stage, and selection intensity on short-term genetic gain. The Plant Genome page e20327

Zhong S, & Jannink J L (2007), Using quantitative trait loci results to discriminate among crosses on the basis of their progeny mean and variance. Genetics 177(1):567–576

